# GUIDES: sgRNA design for loss-of-function screens

**DOI:** 10.1101/168849

**Authors:** Joshua A. Meier, Feng Zhang, Neville E. Sanjana

## Abstract

GUIDES (Graphical User Interface for DNA Editing Screens) is a web-based tool for the design of custom, large-scale CRISPR libraries for loss-of-function screens in human and mouse. GUIDES combines multi-tissue RNA-sequencing data to target expressed exons, protein annotation to target functional domains, sophisticated on-target and off-target guide RNA scoring and other optimizations to create CRISPR libraries directly from a list of genes without requiring any programming expertise.

---

Genome-scale CRISPR-Cas9 knockout libraries have emerged as powerful tools for unbiased, phenotypic screening^1^. These libraries contain a fixed number of Cas9 single-guide RNAs (sgRNAs) targeting each gene in the genome and typically require large numbers of cells (>10^8^) to maintain genome-scale representation. However, there are many applications where it would be preferable to design a custom library targeting specific genes sets (e.g. kinases, transcription factors, chromatin modifiers, the druggable genome) with higher coverage for these specific genes. To address this need, we developed Graphical User Interface for DNA Editing Screens (GUIDES), a web application that designs CRISPR knock-out libraries to target custom subsets of genes in the human or mouse genome (available at http://guides.sanjanalab.org/).

After providing a list of genes (as gene symbols, Ensembl IDs, or Entrez IDs), GUIDES creates a library with multiple sgRNAs to target each gene (**Fig. 1a**). To pick optimal sgRNAs, GUIDES integrates tissue-specific RNA expression, protein structure prediction, Cas9 off-target prediction/avoidance, and Cas9 on-target local sequence preferences in an integrated multi-stage pipeline (**Fig. 1b**).

**Figure 1.**
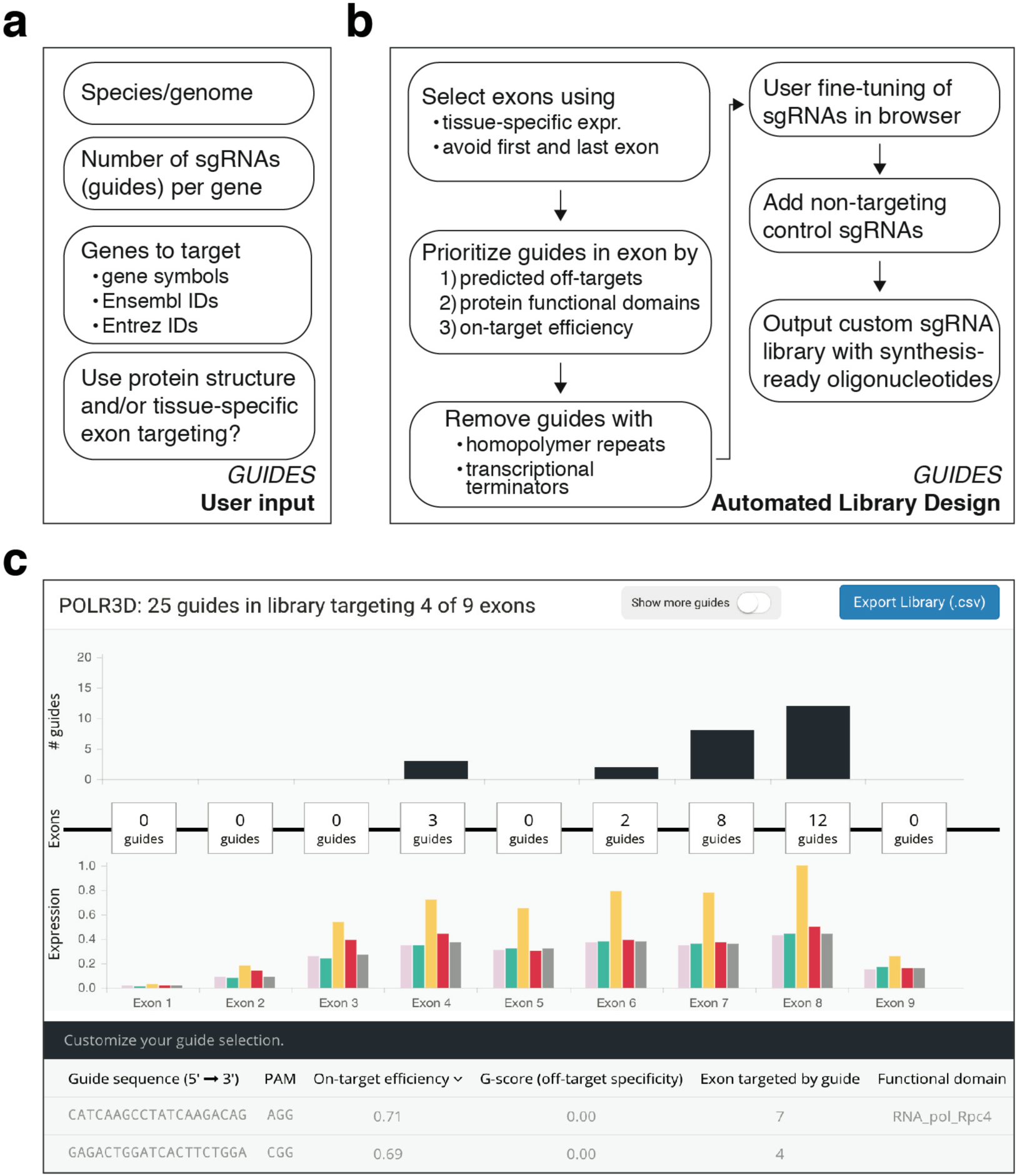
| GUIDES input, workflow and graphical design environment. **(A)** User input for GUIDES automated CRISPR library design. **(B)** Flowchart of GUIDES algorithmic optimization for sgRNA selection. **(C)** Screenshot of interactive designer for adding and deleting genes and sgRNAs. For each sgRNA, the tool displays the exon targeted, the functional domain(s) in the protein targeted (if any), the efficiency (on-target) score, and specificity (off-target) score. The three-panel user interface design allows rapid navigation between gene-level and sgRNA-level changes.

For each gene, GUIDES first identifies coding regions using the Consensus CoDing Sequence (CCDS) database. For the human genome, GUIDES can use tissue-specific RNA-sequencing gene expression data from the GTEx Consortium (v6, 8,555 tissue samples from 544 donors) to target Cas9 preferentially to exons with higher expression^2^. The user can specify specific tissues or compute average expression over a set of tissues (or all tissues), which results in increased targeting to expressed exons (**Supplementary Fig. 1**). Targeting sgRNAs to exons constitutively expressed in the target cell type/tissue can be important, as mutations in alternatively-spliced exons may not result in protein knock-out^3^.

For each exon, GUIDES first prioritizes potential Cas9 target sites using the established cutting frequency determination (CFD) score^4^. For each 20 bp target site, GUIDES identifies all sequences with 1, 2 or 3 base mismatches present in the exome and assigns an off-target score by taking the sum of CFD scores for each potential off-target found. Any target sites with perfect matches elsewhere in the exome are given the maximial off-target score, so that they are selected only in cases where no other sgRNAs exist to target the gene. By using exome-wide CFD scoring during design of a library with ~2,000 genes, the percentage of designed sgRNAs with predicted off-targets decreases from ~43% to ~4% (**Supplementary Fig. 2**).

Although indel mutations are introduced after Cas9 cutting and genome repair, not all indel mutations abolish protein expression, presumably due to in-frame mutations in regions tolerant to mutations. Saturation mutagenesis screens tiling over entire genes have shown increased knock-out efficiency when targeting protein functional domains^5^. To take advantage of this, GUIDES includes an option to preferentially choose sgRNAs that target functional protein domains identified in the Protein Family (Pfam) database (v30, 16,306 protein families)^6^. This can have a significant impact on library design since 90% of protein-coding genes in the human genome contain at least one Pfam-annotated domain^7^.

After identifying exons and protein domains to target, GUIDES uses a previously validated boosted regression tree classifier to score Cas9 target sites based on local sequence preferences learned from saturation mutagenesis screens and adds the highest-scoring sgRNAs to the library^4^. When targeting the same sets of genes, sgRNAs designed with this criterion have a ~30% higher on-target efficiency score (**Supplementary Fig. 3**). Other CRISPR library design tools have also used on-target efficiency scoring to help automate sgRNA design, but do not include RNA expression or protein domain identification to sgRNA targeting^8^ (**Table 1**). Some of the existing library design tools are command-line programs, whereas GUIDES is a graphical, web-based tool that allows fine-tuning of sgRNA selection directly in the web browser (**Fig. 1c**).

**Table 1:**
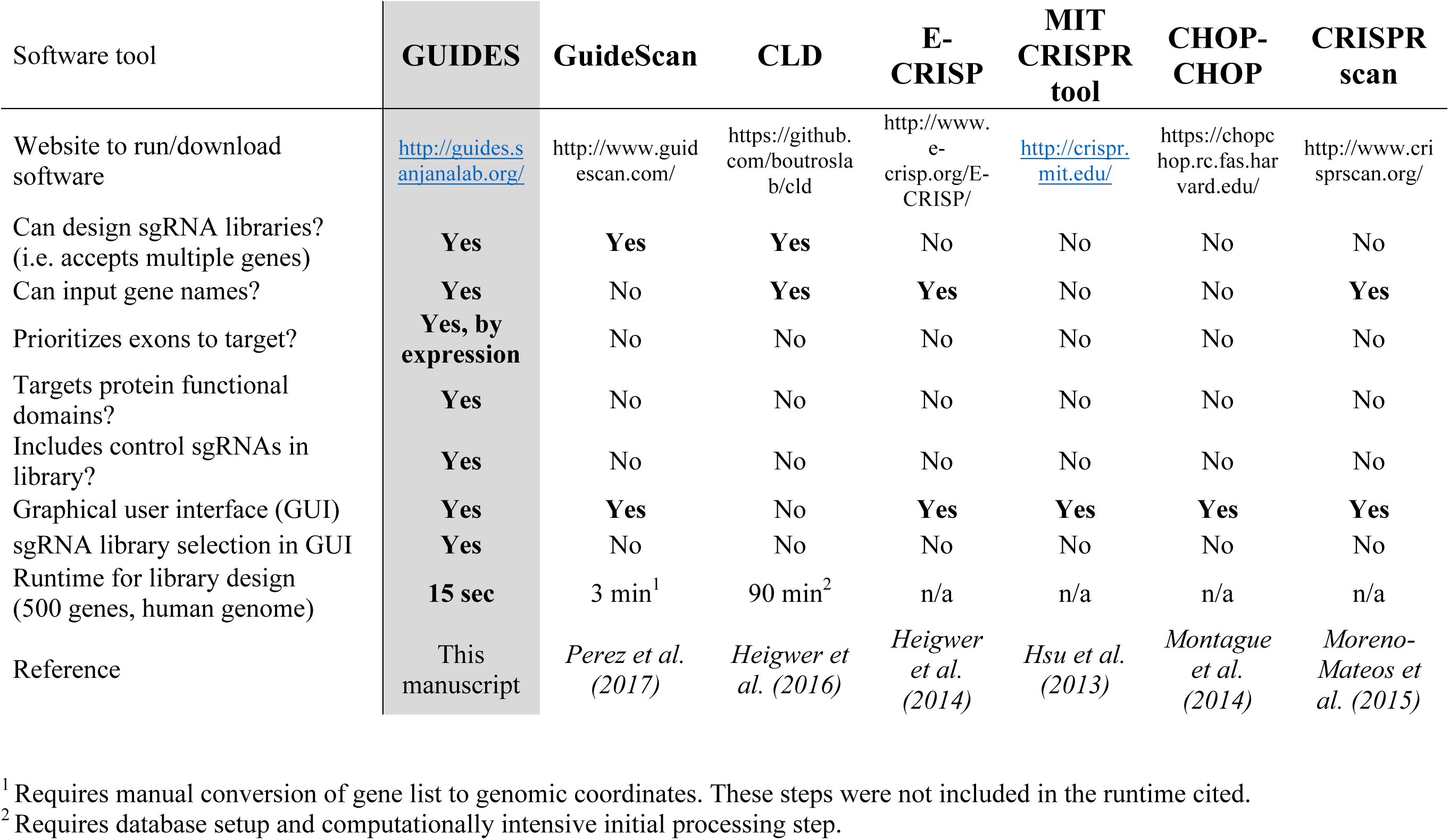
Comparison of CRISPR sgRNA library design tools.

To benchmark the performance of GUIDES-selected sgRNAs in genome-scale screens, we tested whether sgRNAs designed by GUIDES have consistently higher/lower activity using a metaanalysis of 77 pooled CRISPR screens from the GenomeCRISPR database^9^. By examining sgRNAs targeting essential genes, we found that GUIDES-generated sgRNAs were more depleted by approximately one 10%-quantile (with sgRNAs given a percentage rank within each pooled screen) than a size-matched control set of sgRNAs targeting the same gene (**Supplementary Fig. 4**) (*n* = 403 genes with 8 ± 6 sgRNAs per gene, *p* = 5 × 10^−7^, *t* = -5.1, *df* = 409, two-sample paired t-test).

In addition to target site selection, GUIDES also manages several practical aspects of library design, including eliminating sgRNAs with homopolymer repeats that are difficult to synthesize, alerting the user when the sgRNA targets the last exon which may escape nonsense-mediate decay of mRNA^10^, eliminating sgRNAs with Pol3 transcriptional terminators, creating synthesis-ready oligonucleotides with flanking sequences for PCR-based cloning and adding in nontargeting sgRNAs to serve as negative controls. These negative controls can be used to calculate an empirical measure of false-discovery rates in pooled CRISPR screens^11^.

Finally, GUIDES uses several algorithmic optimizations so that, despite the sophisticated design criteria, the user wait times are minimal (**Supplementary Methods**). Run time is linear with respect to gene count and constant with respect to sgRNAs per gene (**Supplementary Fig. 5**). For example, GUIDES takes ~15 seconds to design a library targeting 500 genes involved in chromatin regulation with 6 sgRNAs per gene (Intel i7 3Ghz, 16 GB RAM).

The code for GUIDES and all functional tests is available at https://github.com/sanjanalab/GUIDES. The ready-for-use web tool is available at http://guides.sanjanalab.org/. We hope that GUIDES will allow a wider audience access to custom CRISPR libraries for future loss-of-function forward genetic screens.

## Acknowledgements

We would like to thank B. Cummings for help with GTEx data processing and the entire Zhang and Sanjana laboratories for support and advice. F.Z. is supported by the NIH through NIMH (5DP1-MH100706 and 1R01-MH110049); NSF; the New York Stem Cell Foundation; the Allen Distinguished Investigator Program, through The Paul G. Allen Frontiers Group; the Simons and Vallee Foundations; the Howard Hughes Medical Institute; the Skoltech-MIT Next Generation Program; James and Patricia Poitras and the Poitras Center for Affective Disorders; Robert Metcalfe; and David Cheng. F.Z. is a New York Stem Cell Foundation-Robertson Investigator. N.E.S. is supported by the NIH through NHGRI (R00-HG008171) and a Sidney Kimmel Scholar Award.

## Author contributions

N.E.S. conceived of the library design tool. J.A.M. wrote the code and performed the experiments. N.E.S. and J.A.M. analyzed the experiments. J.A.M., F.Z., and N.E.S. wrote the manuscript. N.E.S. and F.Z. supervised the work.

## Competing financial interests

A patent has been filed relating to the described work. F.Z. is a founder and scientific advisor for Editas Medicine, and a scientific advisor for Horizon Discovery.

